# Improved characterization of single-cell RNA-seq libraries with paired-end avidity sequencing

**DOI:** 10.1101/2024.07.10.602909

**Authors:** John T. Chamberlin, Austin E. Gillen, Aaron R. Quinlan

## Abstract

Prevailing poly(dT)-primed 3’ single-cell RNA-seq protocols generate barcoded cDNA fragments containing the reverse transcriptase priming site, which is expected to be the poly(A) tail or a genomic adenine homopolymer. Direct sequencing across this priming site was historically difficult because of DNA sequencing errors induced by the homopolymeric primer at the ‘barcode’ end. Here, we evaluate the capability of “avidity base chemistry” DNA sequencing from Element Biosciences to sequence through this homopolymer accurately, and the impact of the additional cDNA sequence on read alignment and precise quantification of polyadenylation site usage. We find that the Element Aviti instrument sequences through the thymine homopolymer into the subsequent cDNA sequence without detectable loss of accuracy. The resulting paired-end alignments enable direct and independent assignment of reads to polyadenylation sites, which bypasses complexities and limitations of conventional approaches but does not consistently improve read mapping rates compared to single-end alignment. We also characterize low-level artifacts and arrive at an adjusted adapter trimming and alignment workflow that significantly improves the alignment of sequence data from Element and Illumina, particularly in the context of extended read lengths. Our analyses confirm that Element avidity sequencing is an effective alternative to Illumina sequencing for standard single-cell RNA-seq, particularly for polyadenylation site analyses but do not rule out the potential for similar performance from other emerging platforms.

## Background

Avidity sequencing is a short-read DNA sequencing chemistry and platform recently released by Element Biosciences(Arslan et al., 2023). One purported advantage compared to sequencing-by-synthesis approaches like Illumina is a greatly reduced error rate following homopolymers, as demonstrated in whole genome DNA sequencing(Arslan et al., 2023; Carroll et al., 2023). However, this capability has not been utilized in other contexts. For example, homopolymers occur for technical reasons in many single-cell RNA-seq protocols. In 10x Genomics Chromium 3’ scRNA-seq libraries, a 30bp poly(dT) primer separates the 28bp molecular barcodes from the presumed end of the mRNA, i.e. the polyadenylation site(Zheng et al., 2017). Because the mate pair (reverse) reads typically do not span the entire cDNA insert, sequencing through the primer with a longer forward read should, in principle, simplify the assignment of reads to polyadenylation sites or discrimination of reverse transcriptase artifacts. Due to the sharp reduction in post-homopolymer accuracy of the Illumina technology, this is traditionally avoided, and the commercial Cell Ranger software is hard-coded to ignore sequences after the barcodes. Instead, the ‘standard’ sequencing configuration uses a short, forward read that covers only the barcodes (e.g., 28bp), and a ∼90bp reverse read capturing part, but rarely all, of the cDNA insert fragment (**Figure 1A**). The narrow coverage window impedes the analysis of alternative splicing and isoform diversity(Arzalluz-Luque & Conesa, 2018) and is also sensitive to annotation quality(Cheung et al., 2018; Soneson et al., 2019) or gene-specific mappability(Srivastava et al., 2019). This leaves alternative polyadenylation as the most facile variety of isoform-level analysis, and at least 16 utilities have been released for this task(Ye et al., 2023). Contemporary approaches are limited by two factors: first, most cDNA inserts are longer than the read length; hence, they cannot provide direct evidence of the read’s precise origin. Second, a substantial fraction of cDNAs arise from internal priming rather than poly(A) tails, which requires heuristics to discard potential artifactual sites.

**Figure 1.**
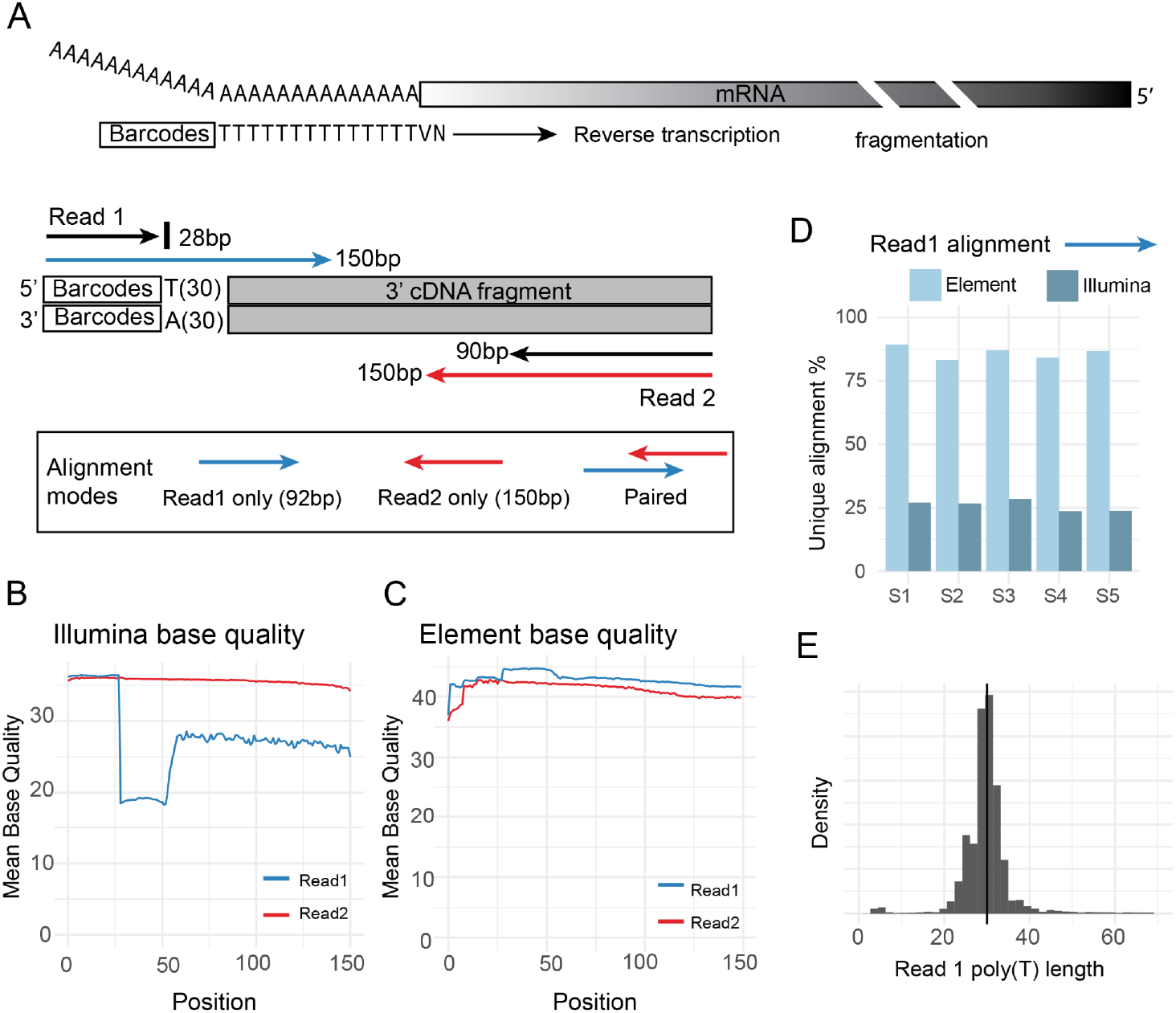
Assessment of Aviti post-homopolymer sequencing accuracy in single-cell RNA-seq. (**A**) Library schematic and sequencing configuration. Barcoded primers are designed to initiate reverse transcription at the base of the poly(T) tail, as primers end in a VN dinucleotide ‘anchor’ sequence (VN = A,C, or G followed by A,T,C, or G) (**B**) Fastq base quality trace for Illumina 150bp x 150bp data, sample S1. Nucleotide positions are reported from 5’ to 3’ for read 1 and read 2. (**C**) Element 150bp x 150bp on the same library reports no loss of quality in the homopolymer segment (**D**) Alignment of Read1 cDNA segment (position 59-150) is reliable with Element but not Illumina sequencing. (**E**) Measured primer (poly(T) homopolymer starting at position 29) has a median length of 30bp, but modest contraction and expansion.

Here, we evaluate the potential for Avidity sequencing to bypass these complications by directly sequencing the end of each cDNA fragment with an extended forward read. We report a simple benchmark using paired-end data from five human single-cell libraries sequenced on the Element Aviti instrument and the Illumina Novaseq6000. We evaluate the merits of paired and single-end alignments and the impact of read length on mappability and the assignment of reads to polyadenylation sites. We also identify necessary adjustments to the alignment workflow to achieve maximal performance and discuss pertinent unrecognized artifacts and their broader relevance. In light of cost considerations amid renewed competition in short-read DNA sequencing, we anticipate that these results and data will be useful to the design and analysis of future experiments.

## Results and Discussion

### Avidity sequencing is not affected by technical homopolymers

We resequenced five human single-cell cDNA libraries from the 10x Genomics 3’ V3 protocol in 150bp x 150bp configuration using the Element Aviti instrument, generating about 1.8 billion reads from two flowcells. Each library was previously sequenced on an Illumina Novaseq 6000 in the same configuration (**Figure 1A**)(Pei et al., 2020, 2023). As expected, Illumina forward reads (R1) exhibited a marked drop in average base quality in the defined homopolymer interval (position 29-58) and only partial recovery after emerging into the mRNA segment (**Figure 1B**). In contrast, Element data exhibited a high base quality across the length of both reads and a mild increase in base quality in the homopolymer window in R1 (**Figure 1C**), which may indicate base quality miscalibration. The post-homopolymer performance is consistent with previous data from PCR-free whole-genome sequencing(Arslan et al., 2023).

We validated the instrument-reported base qualities by independently aligning the cDNA interval of the forward reads (position 59-150 of R1) in single-end mode (**Figure 1D**). Element data showed unique mapping rates of 80-88%, which is similar to the threshold described by 10x Genomics(*Quality Assessment Using the Cell Ranger Web Summary*, n.d.). Mismatch rates per base pair were less than 0.5%. Illumina R1 mapping rates hovered around 25%, and mismatch rates per base pair were above 5%. The median poly(T) homopolymer length observed in Element R1 was 30bp, as expected, but with modest variation (**Figure 1E**): the middle 90% of reads showed poly-T lengths between 23bp and 38bp. These results demonstrate that Aviti’s homopolymer accuracy also translates to scRNA-seq analysis. However, the variation in measured homopolymer length suggests some expansion and contraction of the primer during PCR or sequencing, and that adjustments to the bioinformatic workflow are needed to maximize read alignment and resolution of polyadenylation sites.

### After necessary workflow adjustments, paired-alignment reduces multi-mapping but not overall alignment rate

We next evaluated the impact of paired-end sequences on read alignment in Element data. We expected paired alignment to reduce multimapping, and R2 alignments to be better than R1 alignments due to the differences in effective read length (150bp vs 92bp). Instead, we initially found that unique *paired* alignment was higher than *R2*, but not always higher than *R1*, with substantial variation across libraries. Investigation of mapping statistics and unaligned reads suggested that increased adapter and primer content in the finished libraries was likely responsible. Adapters or primers are expected, particularly with increased read lengths, but we observed substantial variation across libraries, particularly in the level of poly(A) read-in (**Figure 2A**). Adapter content was similar between Illumina and Element data, indicating the variation is a result of sample variability at the library preparation stage. While STAR includes a built-in adapter trimming feature (modeled after the same functionality in Cell Ranger), it is not configured for paired-end sequences, and we found the impact on single-end alignments to be small (**Figure 2B**).

**Figure 2.**
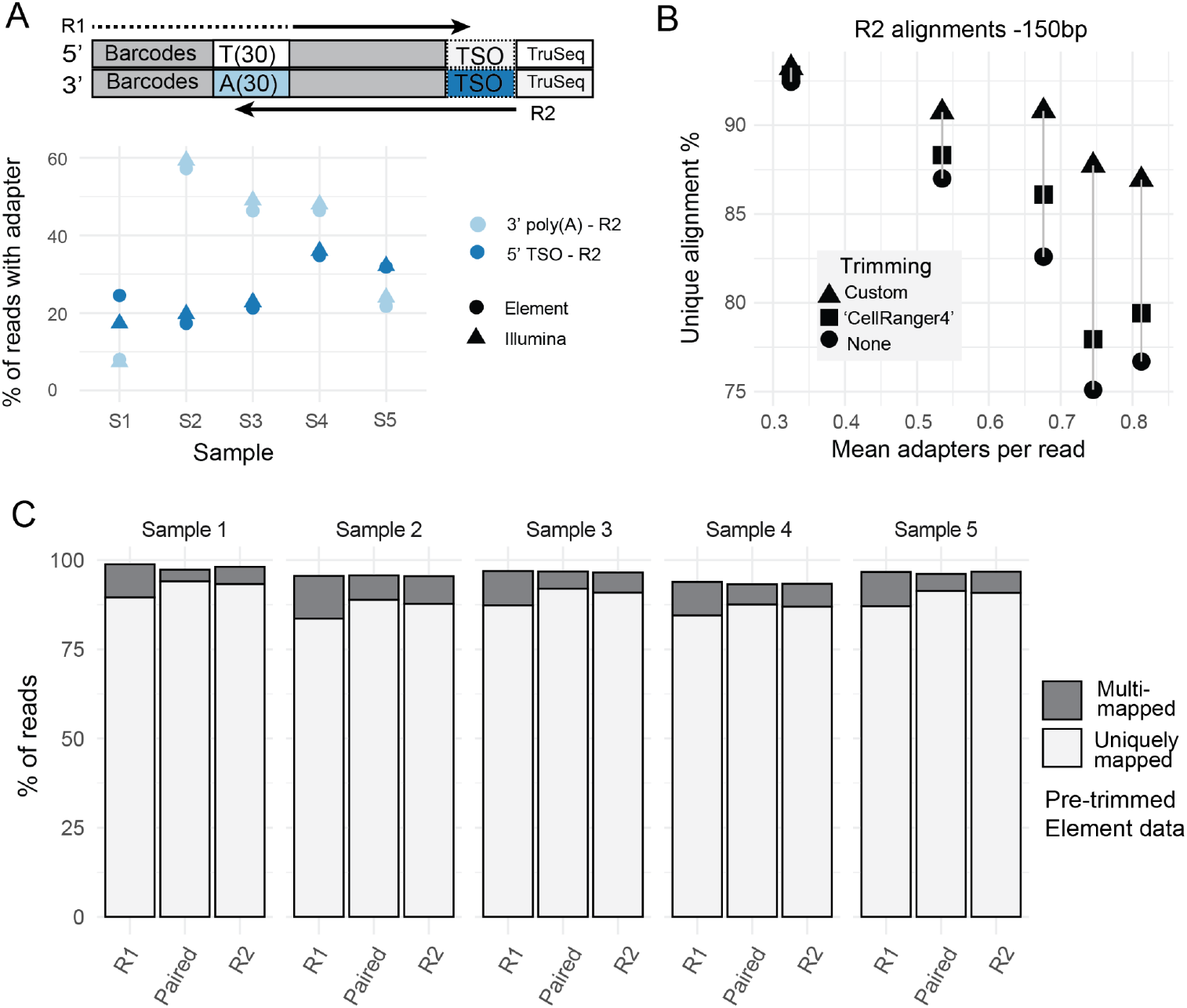
Extended and paired-end reads require additional preprocessing. (**A**) Adapter schematic in finished libraries. 3’ poly(A) will appear in any R2 longer than the cDNA insert. Most 5’ TSO primers are lost during cDNA fragmentation step, but will be sequenced at the beginning of R2 if they remain in the finished library. Adapter content (reads with one or more adapters) is consistent between sequencers but highly variable across samples, indicating variation in cDNA insert size distribution. (**B**) Adapter pre-trimming improves single-end unique alignment rate of 150bp R2 compared to the current implementation of Cell Ranger or STARsolo, particularly in samples with higher adapter content. Unique alignment rate declines according to adapter content regardless of trimming, likely due to the occurrence of reads with very short cDNA inserts. ‘Custom’ trimming metrics account for reads discarded prior to alignment based on a minimum length threshold of 31bp. Mean adapters per read refers to the total number of trimmed TSO and p(A) sequences divided by the total number of reads. A single read may contain 0, 1, or 2 adapters. (**C**) Even after optimized trimming and alignment parameterization, paired-end alignment slightly improves the unique alignment rate but not the overall alignment rate (unique + multi-mapped reads) compared to R2 or R1 single-end alignment. R1 alignments show considerably higher rates of multi-mapping due to shorter effective read lengths, but no elevation in mismatch rate at the per-nucleotide level.

As such, we adopted an adapter pre-trimming approach (**Methods**) which removes 5’ and 3’ primers and adapters from both mate pairs. This approach substantially increased R2 alignment rates, particularly in samples with higher adapter content (**Figure 2B**). Incidentally, we found that the built-in adapter trimming was much more effective if R2 was first hard-clipped to 90bp, suggesting that it fails to account for non-terminal poly(A) homopolymers, which will occur if R2 fully traverses the primer and enters the barcode segment. We repeated the alignment benchmark for R1, paired, and R2 alignment on the same pre-trimmed data; paired-end unique alignment rates ranged from 87.5% to 94%, marginally higher than R2 single-end alignment. Nevertheless, we found that the overall mapping rate (including multi-mapped reads) was not consistently better for paired vs single-end alignments (**Figure 2C**). Instead, longer effective read lengths (paired > R2 > R1) shifted mapping profiles from multi-to uniquely mapped. Multi-mapping was ∼1.6-2.9X higher for R1 than paired alignment. Most of the ‘unmapped’ reads were the result of up to 4% of reads falling below the trimmed read length threshold, indicative of extremely short cDNA inserts (31bp, see **Methods**). Of the reads passed to the aligner, 1.5% to 2.5% failed to align. Between technologies, Illumina R2 alignment performed well, but unique alignment rates were 2-7 percentage points lower than Element (**Supplementary Table 1**). However, we can not discount the possibility that homopolymer sequencing in Illumina R1 has a carryover effect on base calling in R2. We also compared gene expression estimates between technologies but found them to be highly consistent (R > 0.92, **Supplementary Table 2**).

These results suggest that while Avidity sequencing generates highly accurate data from single-cell RNA-seq libraries, the homopolymer performance is not inherently advantageous for gene-level single-cell RNA-seq analysis. However, paired alignments do allow direct estimation of cDNA insert size distribution. Variation across our samples suggests caution is warranted when attempting to make use of size distribution inferred from independent samples, as was done in a recent preprint to improve discrimination of spliced and unspliced transcripts (He et al., 2024). Finally, we note that the adapter content metrics shown here were manually collated from the output of the adapter pre-trimming step; these values are not reported by typical pre-processing workflows such as STARsolo or Cell Ranger. Similarly, the basis for read misalignment (i.e., insufficient alignment length) is not reported by Cell Ranger. Hence, the observed detrimental effect of increased adapter read-in with 150bp R2 length is a useful and unforeseen byproduct of our investigation into paired-end alignment performance.

### Paired-end alignments improve the resolution of polyadenylation sites

At least 16 methods (**Supplementary Table 3**) have been developed for quantifying polyadenylation site usage from traditional single-end (R2) scRNA-seq alignments(Ye et al., 2023). While a comprehensive benchmark has not been performed, the adoption of these tools in biological studies also remains somewhat limited. We selected polyApipe to test the utility of the traditional single-end alignment approach, including after hard-clipping R2 to 90bp. While unpublished, this tool was used by a recent publication on polyadenylation machinery(Kowalski, 2024) and has also been incorporated into the popular Seurat suite of R packages for single-cell analysis. More importantly, it is simple in principle and thus serves as a useful and intuitive comparator for the broader discussion of paired and single-end alignments. PolyAPipe relies on soft-clipped strings of A’s at the end of R2 alignments to identify putative polyadenylation sites and assigns additional reads to these sites if they align in the upstream 250bp (**Figure 3A**). It then flags presumed off-target sites (internal priming sites) based on the presence of an arbitrarily defined genomic poly(A) homopolymer. This approach avoids complexities in other workflows such as peak-calling or size distribution modeling, but may also lose reads that align across splice junctions, mis-aggregate distinct polyadenylation sites into a single ‘peak’, or mistake true sites for internal priming sites or vice versa. Reliance on soft-clipping also sensitizes the approach to variation in insert size distribution or shortened read lengths, as only the cDNAs with inserts shorter than the read length can exhibit the soft-clipped signal.

**Figure 3.**
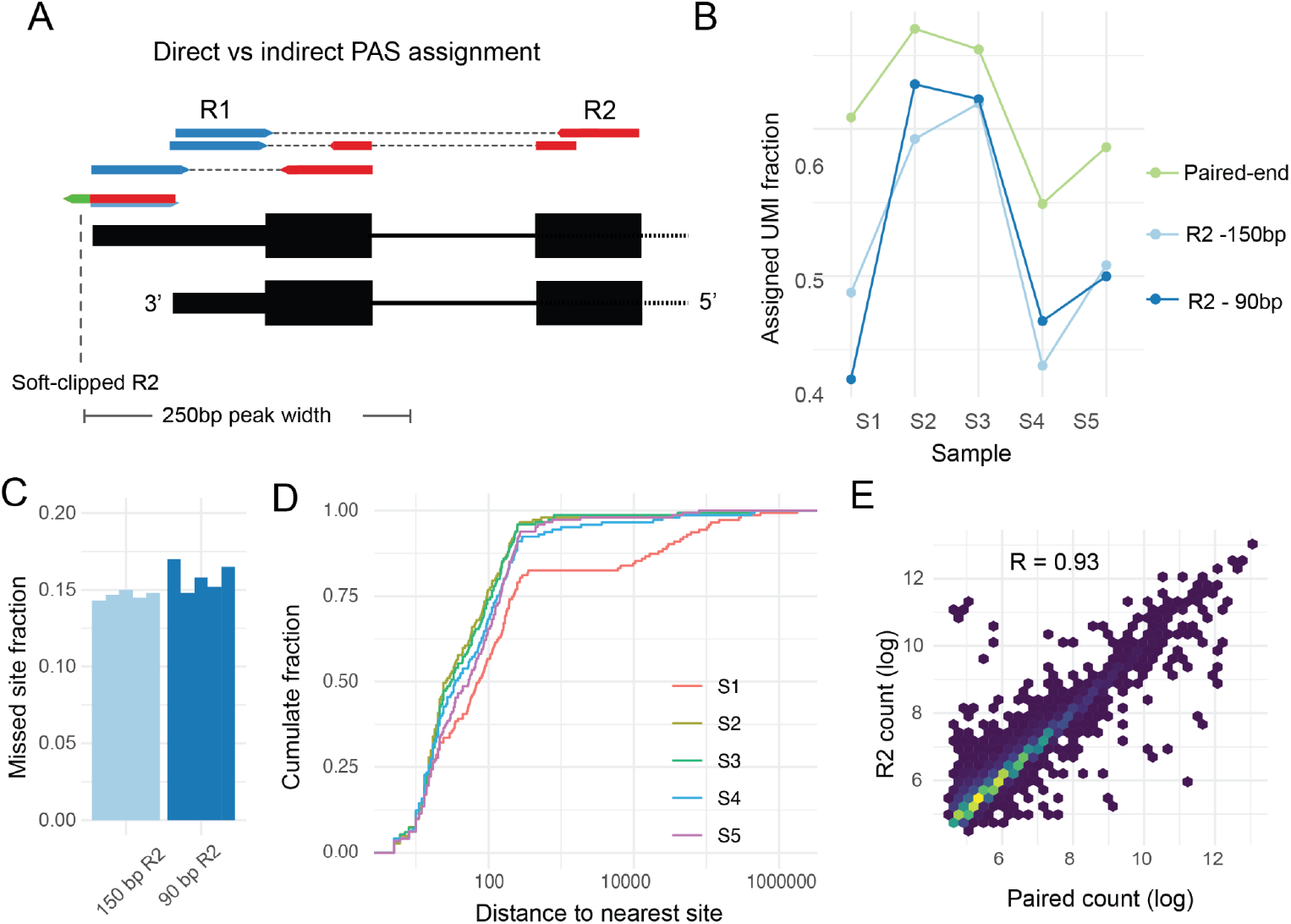
Impact of paired-end alignments of polyadenylation site analysis. (**A**) Schematic describing traditional single-end approach vs paired-end approach. PolyApipe looks for soft-clipped A’s at the 3’ end of R2 alignments (green) and then allocates additional R2 alignments within a definable window. With the addition of R1 alignments, scraps directly assigns alignments based on the alignment start coordinate of R1. Note that the gene model is depicted in 3’ to 5’ orientation (**B**) Paired-end assignment increases the overall rate at which exonic transcripts can be assigned to annotated polyadenylation sites. 90bp R2 outperforms some 150bp R2 for some libraries due to improved alignment rate from builtin adapter trimming mode. PolyApipe ‘misprime’ peaks are not excluded. (**C**) Single-end assignment misses a substantial fraction of the top 1000 sites detected from paired-end alignments, based on a window of -10/+5 bp around annotated sites. (**D**) Of the top 1000 paired-end sites missed by single-end assignment, most are within 100bp of an alternative single-end assigned site. The median distance ranges from 26 to 73bp across samples, varying according to library insert size (**E**) Total scraps*-*assigned UMIs vs total polyApipe-assigned UMIs at shared sites in Element library S1. Log-transformed counts are highly correlated between paired and single-end assignments, R=0.93. Correlation across remaining samples was comparable, R=0.99.

For paired-end alignment, we used the scraps workflow(Fu et al., 2022), which quantifies polyadenylation sites defined in the PolyADB_V3 set of annotated polyadenylation sites; these sites were defined using the 3’READS assay which is designed to avoid internal priming artifacts.(Wang et al., 2018). By default, scraps assigns reads based on the 5’ start of the R1 alignment (**Figure 3A**), allowing for -10/+5bp buffer. We reduced the R1 soft clipping parameter of the R1 barcode/primer segment in the alignment step from 58bp to 48bp (**Methods**) to account for the contraction of the primer sequence, which would create a spurious offset in the apparent polyadenylation site (**Figure 1E**). We measured overall polyadenylation site assignment performance as the fraction of exonic transcripts (i.e. UMIs, or Unique Molecular Identifiers) assigned by either workflow to an externally-annotated polyadenylation site. This step is integral to the scraps workflow, while for polyApipe we intersected the 3’ coordinate of each ‘peak’ with the same set of annotated polyadenylation sites (again allowing -10/+5bp). We found that scraps assigned between 55 and 67% of exonic UMIs to annotated polyadenylation sites, while polyApipe assigned between 44 and 62% with 150bp R2 and between 43 and 63% with 90bp R2 (**Figure 3B**). The inconsistent effect of read length on polyApipe is due to read alignment effects, as the adapter pre-trimming step is not compatible with the search for soft-clipped tails. PolyApipe flagged about 20% of its peaks as ‘misprimed’, but we did not discard these given that they intersected with annotated polyadenylation sites. We also observed discrepant behavior in the misprime-calling step, as more sites were flagged with 90bp than with 150bp R2. The reduction in UMI assignment is largely driven by highly-expressed polyadenylation sites that are absent from the polyApipe output: about 15% of the 1000 most abundant *scraps*-assigned polyadenylation sites were not ‘detected’ by polyApipe, increasing slightly with 90bp R2 (**Figure 3C**). This indicates that the single-end reads in this vicinity, while presumably still present, were assigned to an alternative ‘peak’ boundary more than -10/+5 away from the annotated polyadenylation site. Differences between 150bp and 90bp R2 is consistent with the library size variation: 5%-55% 150bp R2 contained a poly(A) primer (**Figure 2A**) vs just 1-12% of 90bp reads, with the biggest difference in samples S1 and S5.

For the sites that were identified by scraps but not by polyApipe, about 50% were at least 40bp away from the nearest polyApipe-defined site, again varying with insert size distribution (**Figure 3D**). However, transcript counts were highly correlated at shared sites (**Figure 3E**). These patterns support the premise that paired alignments improve the resolution of polyadenylation sites, even at highly-expressed sites, but also indicate that the net benefit is relatively small. Finally, we manually inspected outlier sites to substantiate the basis for discrepancies between the two approaches. For example, *COX6C* is detected almost exclusively from an isoform with a short terminal exon (145bp). In libraries with shorter insert sizes (S2,3 and 4), paired and single-end quantification yield comparable counts. However, library S1, with its longer insert sizes, shows a 20-fold disparity due to R2 alignments shifting into the adjacent exon, outside the 250bp window used by polyApipe (**Supplementary Figure 1**).

Conversely, the polyadenylation site for *SOD2* is 40-fold more prevalent in the R2 polyApipe result than in the paired-end scraps result. Inspection of alignments shows this is because the gene is almost exclusively utilizing an unannotated polyadenylation site 51bp upstream of the annotated coordinate (**Supplementary Figure 2**). PolyApipe allocates all of these reads to the distal site, while scraps omits them. This case highlights the limitation of relying on a database of annotated sites for the current implementation of scraps. However, it clearly demonstrates the potential utility of direct sequencing of the polyadenylation site in the context of closely-spaced alternative polyadenylation - about 30% of polyadenylation sites in our annotation are within 100bp of another polyadenylation site. These results confirm that paired-end avidity sequencing improves base-pair precision of polyadenylation site assignment, but the benefit is largely restricted to the subset of challenging regions characterized by multiple nearby polyadenylation sites or polyadenylation sites very close to splice junctions.

## Conclusion

We validated the homopolymer performance of Avidity sequencing from Element Biosciences by applying it to paired-end sequencing of 3’ single-cell RNA-seq libraries. We show that sequencing across the reverse transcriptase priming site in each read pair enables independent and accurate resolution of polyadenylation site usage and reduced variance induced by sample-specific insert size distribution. However, paired-end sequences have a minimal impact on read alignability itself, evidently due to the overall short insert size distribution favored by the single-cell protocol. Instead, insert size distribution appears to be a stronger contributor to the upper limit of read alignment performance. We observed tradeoffs among our samples between insert size distribution, alignment, and polyadenylation site analysis: longer inserts improve alignment and reduce adverse effects of primer read-in, but also worsen polyadenylation site assignment performance for single-end methods which rely on soft-clipping. Adjustments to the size selection step could make better use of this capability. We used a symmetrical sequencing configuration out of simplicity. However, future studies may find it preferable to allocate instrument cycles in an asymmetric or even one-sided manner. For example, the Ultima genomics platform relies on longer single-end reads, and has also shown utility for single-cell RNA-seq(Simmons et al., 2023). In summary, our study provides useful guidance for the design and analysis of future single-cell RNA-seq studies amid the emergence of competing short-read sequencing platforms and addresses important informatics impediments to the handling of paired-end or extended read-length data with conventional tools. The utility of Avidity sequencing for other assays in which homopolymers are present also warrants further consideration. The data generated here are made publicly available to support future bioinformatic methods development.

## Methods

### Single-cell library preparation and sequencing

The initial sample dissociation and library preparation for Illumina sequencing are described in two previous publications(Pei et al., 2020, 2023). All samples used the 10x Genomics V3 protocol, i.e., 12bp UMI and 16bp cell barcode. Sample ‘S1’ specifically used v3.0 while the other four libraries used v3.1. Illumina sequencing was performed on a NovaSeq 6000 instrument in the Genomics Shared Resource at the University of Colorado Cancer Center. Excess material from finished libraries was later shipped to the University of Utah Core where it was prepared for Element sequencing using the ‘Adept’ circularization kit. We sequenced five libraries across two Aviti flow cells, generating about 1.8 billion total read pairs. Sequencing metrics are summarized in **Table S1**.

### Adapter trimming

We used cutadapt(Martin, 2011) and selected utilities from seqtk(Li, n.d.) and bbtools(Bushnell, 2014) to manually trim primers and adapters from fastq files before alignment. Specifically, we trimmed 3’ poly(A) and 5’ TSO from R2, excess 5’ poly(T) from R1 (to account for the variation seen in Figure 1D), and 3’ TSO and TruSeq primers from R2 with cutadapt. We collated adapter trimming stats from the json output of cutadapt. For R2 single-end alignment only, we also tested STAR’s built-in adapter trimming feature ‘--clipAdapterType CellRanger4’ which is meant to trim 3’ poly(A) and 5’ TSO sequences. We used seqtk to hard-clip R2 to 90bp before some alignment tests, and used the ‘repair’ utility from bbtools to discard reads with a missing mate, when necessary.

### Alignment and quantification

All alignment and gene-level transcript quantification used STARsolo version 2.7.9a(Kaminow et al., 2021). To use this version of STAR and its latest features, we re-indexed the 2024-A human GRCh38 reference annotation provided by 10x Genomics, which uses a filtered set of gene and transcript models from GENCODE v44. We used the ‘Gene’ quantification mode in STARsolo to quantify genes based on exonic alignments.

### Alignment and trimming workflow optimization

Because STARsolo is not commonly used in paired-end configuration, we found that certain non-default parameters needed to be enabled. We first activated the ‘peOverlapNbasesMin’ feature which enables merging and ‘second-pass’ alignment of overlapping mate pairs. We also enabled ‘alignEndsProtrude’ which allows for alignments to extend past each other, which might occur due to imbalances in the trimming step. We set a minimum alignment length of 31bp based on the default behavior observed in the output of Cell Ranger v8. This value signifies that any trimmed read which is shorter than 31bp or which yields an aligned subsequence shorter than 31bp will be called as unmapped. We did not adjust the mismatch parameters for the analyses described here; however, a variety of other options are available for relaxing the alignment or mate-pair merging thresholds, which may lead to marginal increases in read alignments.

For single-end alignment, we tested alignment with no adapter trimming, built-in trimming (--clipAdapterType CellRanger4), and with the same pre-trimming used for the paired-end alignments. We also confirmed that Cell Ranger exhibited the same failure to properly trim 150bp R2 reads, i.e. it exhibited a worse alignment rate for 150bp R2 than for hard-clipped 90bp R2.

### Polyadenylation site assignment and quantification

We used scraps for paired-end polyadenylation site quantification(Fu et al., 2022). To our knowledge, this is the only workflow currently available for the quantification of polyadenylation sites from paired-end or read1-only sequences. We utilized polyApipe for single-end polyadenylation site quantification. Both workflows utilize umi-tools internally(Smith et al., 2017). We modified scraps by disabling a step that adjusts alignment positions based on the expected size of the primer based on our observation that reads with expanded or contracted primers still aligned to the same boundary. We also adjusted the preceding alignment step by soft-clipping the first 48bp of R1 rather than the first 58bp to account for cases in which the measured primer length contracts by up to 10bp.

We ran polyApipe on BAM files generated with the STARsolo ‘CellRanger4’ adapter clipping option, which soft-clips poly(A) primers rather than removing them entirely. PolyApipe identifies putative PAS from soft-clipped reads and then aggregates other reads within a 250bp upstream window, by default. It then flags sites that contain an aligned stretch of A’s ahead of the soft-clipped tail, as this is indicative of internal priming. We intersected the resulting windows with the same database of validated PAS used by scraps. The 3’ end of the peak had to intersect with a 15bp window around the annotated site, which is the same logic used by scraps. We did not experiment with other parameters of polyApipe, such as the minimum number of soft-clipped reads per site (default = 1) or length of soft-clipped tail.

## Statistical Analyses

All statistical analyses and figure generation were conducted in RStudio(RStudio, n.d.) 2021.09.0, with R version 4.1.1(R Core Team, n.d.). We made extensive use of tidyverse(Wickham et al., 2019) packages. Interval comparisons were performed with GenomicRanges(Lawrence et al., 2013). Figures were generated with ggplot2(Wickham, 2011).

## Code availability

Scripts for adapter trimming, alignment, and polyadenylation site analysis are available at github.com/johnchamberlin/avidity_scrna_benchmark

## Data availability

Raw Illumina data are available on NCBI GEO from these accessions: SRX7528394, SRX20356215, SRX20356216, SRX20356222, SRX20356230.

Element data will be uploaded to GEO and linked to their respective experiments.

## Acknowledgements

We appreciate the technical expertise of Derek Warner at the Utah DNA Sequencing and Genomic Core Facilities for facilitating the Element sequencing.

## Author Contribution

J.T.C. conceived of the study, performed the primary analyses, and wrote the manuscript. A.E.G. oversaw the development of the scraps workflow and provided the Illumina data. A.R.Q. supervised the analysis. All authors assisted with the study formulation and writing.

## References

Arslan, S., Garcia, F. J., Guo, M., Kellinger, M. W., Kruglyak, S., LeVieux, J. A., Mah, A. H., Wang, H., Zhao, J., Zhou, C., Altomare, A., Bailey, J., Byrne, M. B., Chang, C., Chen, S. X., Cho, B., Dennler, C. N., Dien, V. T., Fuller, D., … Previte, M. (2023). Sequencing by avidity enables high accuracy with low reagent consumption. Nature Biotechnology. 10.1038/s41587-023-01750-7

Arzalluz-Luque, Á., & Conesa, A. (2018). Single-cell RNAseq for the study of isoforms-how is that possible? Genome Biology, 19(1), 110.

Bushnell, B. (2014). BBTools software packag. E . https://cir.nii.ac.jp/crid/1370294643771707027

Carroll, A., Kolesnikov, A., Cook, D. E., Brambrink, L., Wiseman, K. N., Billings, S. M., Kruglyak, S., Lajoie, B. R., Zhao, J., Levy, S. E., McLean, C. Y., Shafin, K., Nattestad, M., & Chang, P.-C. (2023). Accurate human genome analysis with Element Avidity sequencing. In bioRxiv (p. 2023.08.11.553043). 10.1101/2023.08.11.553043

Cheung, L. Y. M., George, A. S., McGee, S. R., Daly, A. Z., Brinkmeier, M. L., Ellsworth, B. S., & Camper, S. A. (2018). Single-Cell RNA Sequencing Reveals Novel Markers of Male Pituitary Stem Cells and Hormone-Producing Cell Types. Endocrinology, 159(12), 3910–3924.

Fu, R., Riemondy, K. A., Sheridan, R. M., Hesselberth, J. R., Jordan, C. T., & Gillen, A. E. (2022). scraps: an end-to-end pipeline for measuring alternative polyadenylation at high resolution using single-cell RNA-seq. In bioRxiv (p. 2022.08.22.504859). 10.1101/2022.08.22.504859

He, D., Gao, Y., Chan, S. S., Quintana-Parrilla, N., & Patro, R. (2024). Forseti: A mechanistic and predictive model of the splicing status of scRNA-seq reads. bioRxiv : The Preprint Server for Biology. 10.1101/2024.02.01.577813

Kaminow, B., Yunusov, D., & Dobin, A. (2021). STARsolo: accurate, fast and versatile mapping/quantification of single-cell and single-nucleus RNA-seq data. In bioRxiv (p. 2021.05.05.442755). 10.1101/2021.05.05.442755

Kowalski, H. (2024). The 2024 Michael Bonfiglio Award for Student Research in Orthopaedic Surgery The 2024 Iowa Orthopaedic Society Medical Student Research Award for Musculoskeletal Research. The Iowa Orthopaedic Journal, 44(1), xvii.

Lawrence, M., Huber, W., Pagès, H., Aboyoun, P., Carlson, M., Gentleman, R., Morgan, M. T., & Carey, V. J. (2013). Software for computing and annotating genomic ranges. PLoS Computational Biology, 9(8), e1003118.

Li, H. (n.d.). seqtk: Toolkit for processing sequences in FASTA/Q formats. Github. Retrieved July 1, 2022, from https://github.com/lh3/seqtk

Martin, M. (2011). Cutadapt removes adapter sequences from high-throughput sequencing reads. EMBnet.journal, 17(1), 10–12.

Pei, S., Pollyea, D. A., Gustafson, A., Stevens, B. M., Minhajuddin, M., Fu, R., Riemondy, K. A., Gillen, A. E., Sheridan, R. M., Kim, J., Costello, J. C., Amaya, M. L., Inguva, A., Winters, A., Ye, H., Krug, A., Jones, C. L., Adane, B., Khan, N., … Jordan, C. T. (2020). Monocytic Subclones Confer Resistance to Venetoclax-Based Therapy in Patients with Acute Myeloid Leukemia. Cancer Discovery, 10(4), 536–551.

Pei, S., Shelton, I. T., Gillen, A. E., Stevens, B. M., Gasparetto, M., Wang, Y., Liu, L., Liu, J., Brunetti, T. M., Engel, K., Staggs, S., Showers, W., Sheth, A. I., Amaya, M. L., Minhajuddin, M., Winters, A., Patel, S. B., Tolison, H., Krug, A. E., … Jordan, C. T. (2023). A Novel Type of Monocytic Leukemia Stem Cell Revealed by the Clinical Use of Venetoclax-Based Therapy. Cancer Discovery, 13(9), 2032–2049.

Quality Assessment Using the Cell Ranger Web Summary. (n.d.). 10x Genomics. Retrieved June 21, 2024, from https://www.10xgenomics.com/analysis-guides/quality-assessment-using-the-cell-ranger-web-summary

R Core Team. (n.d.). R: A Language and Environment for Statistical Computing. R Foundation for Statistical Computing. https://www.R-project.org

RStudio, T. (n.d.). RStudio: integrated development for R. Rstudio Team, PBC, Boston, MA URL http://www.paperpile.com/b/hdMosA/Y6xF

Simmons, S. K., Lithwick-Yanai, G., Adiconis, X., Oberstrass, F., Iremadze, N., Geiger-Schuller, K., Thakore, P. I., Frangieh, C. J., Barad, O., Almogy, G., Rozenblatt-Rosen, O., Regev, A., Lipson, D., & Levin, J. Z. (2023). Mostly natural sequencing-by-synthesis for scRNA-seq using Ultima sequencing. Nature Biotechnology, 41(2), 204–211.

Smith, T., Heger, A., & Sudbery, I. (2017). UMI-tools: modeling sequencing errors in Unique Molecular Identifiers to improve quantification accuracy. Genome Research, 27(3), 491–499.

Soneson, C., Love, M. I., Patro, R., Hussain, S., Malhotra, D., & Robinson, M. D. (2019). A junction coverage compatibility score to quantify the reliability of transcript abundance estimates and annotation catalogs. Life Science Alliance, 2(1). 10.26508/lsa.201800175

Srivastava, A., Malik, L., Smith, T., Sudbery, I., & Patro, R. (2019). Alevin efficiently estimates accurate gene abundances from dscRNA-seq data. Genome Biology, 20(1), 65.

Wang, R., Nambiar, R., Zheng, D., & Tian, B. (2018). PolyA_DB 3 catalogs cleavage and polyadenylation sites identified by deep sequencing in multiple genomes. Nucleic Acids Research, 46(D1), D315–D319.

Wickham, H. (2011). Ggplot2. Wiley Interdisciplinary Reviews. Computational Statistics, 3(2), 180–185.

Wickham, H., Averick, M., Bryan, J., Chang, W., McGowan, L., François, R., Grolemund, G., Hayes, A., Henry, L., Hester, J., Kuhn, M., Pedersen, T., Miller, E., Bache, S., Müller, K., Ooms, J., Robinson, D., Seidel, D., Spinu, V., … Yutani, H. (2019). Welcome to the tidyverse. Journal of Open Source Software, 4(43), 1686.

Ye, W., Lian, Q., Ye, C., & Wu, X. (2023). A Survey on Methods for Predicting Polyadenylation Sites from DNA Sequences, Bulk RNA-seq, and Single-cell RNA-seq. Genomics, Proteomics & Bioinformatics, 21(1), 67–83.

Zheng, G. X. Y., Terry, J. M., Belgrader, P., Ryvkin, P., Bent, Z. W., Wilson, R., Ziraldo, S. B., Wheeler, T. D., McDermott, G. P., Zhu, J., Gregory, M. T., Shuga, J., Montesclaros, L., Underwood, J. G., Masquelier, D. A., Nishimura, S. Y., Schnall-Levin, M., Wyatt, P. W., Hindson, C. M., … Bielas, J. H. (2017). Massively parallel digital transcriptional profiling of single cells. Nature Communications, 8, 14049.

