## SupplementaryMaterials for "Improved characterization of single-cell RNA-seq libraries with paired-end avidity sequencing"

### Supplementary Materials

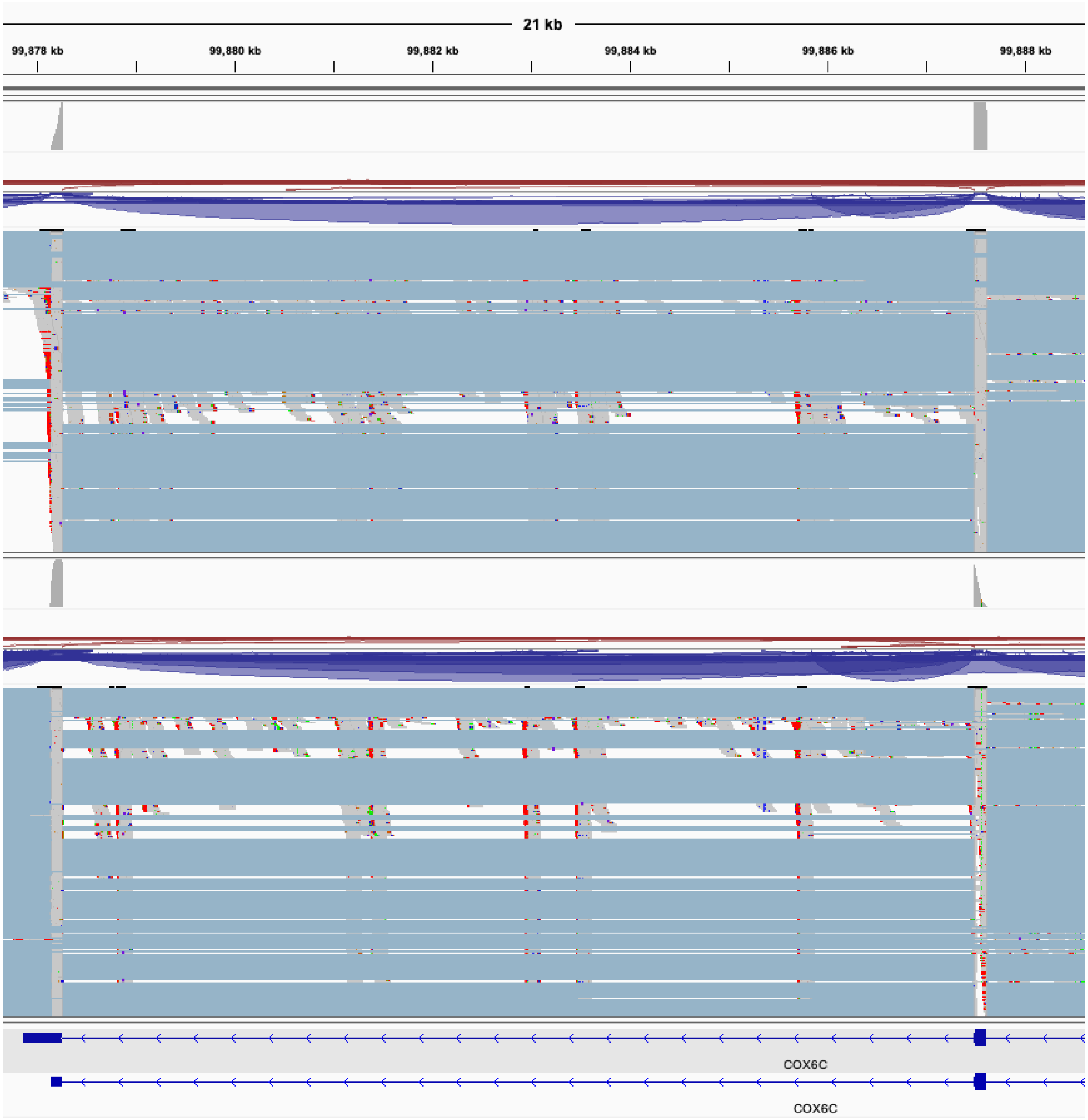


**Supplementary Figure S1**. Single-end read coverage at COX6C exemplifies the impact of short terminal exons. Element library S1, top, has a longer insert size distribution (median 266bp) such that most reads align at the neighboring exon and are missed by the single-end assignment approach. Element library S3, bottom, has a shorter insert size distribution (median 157) such that reads almost always contain the splice junction, leading to similar quantification to paired-end approach.

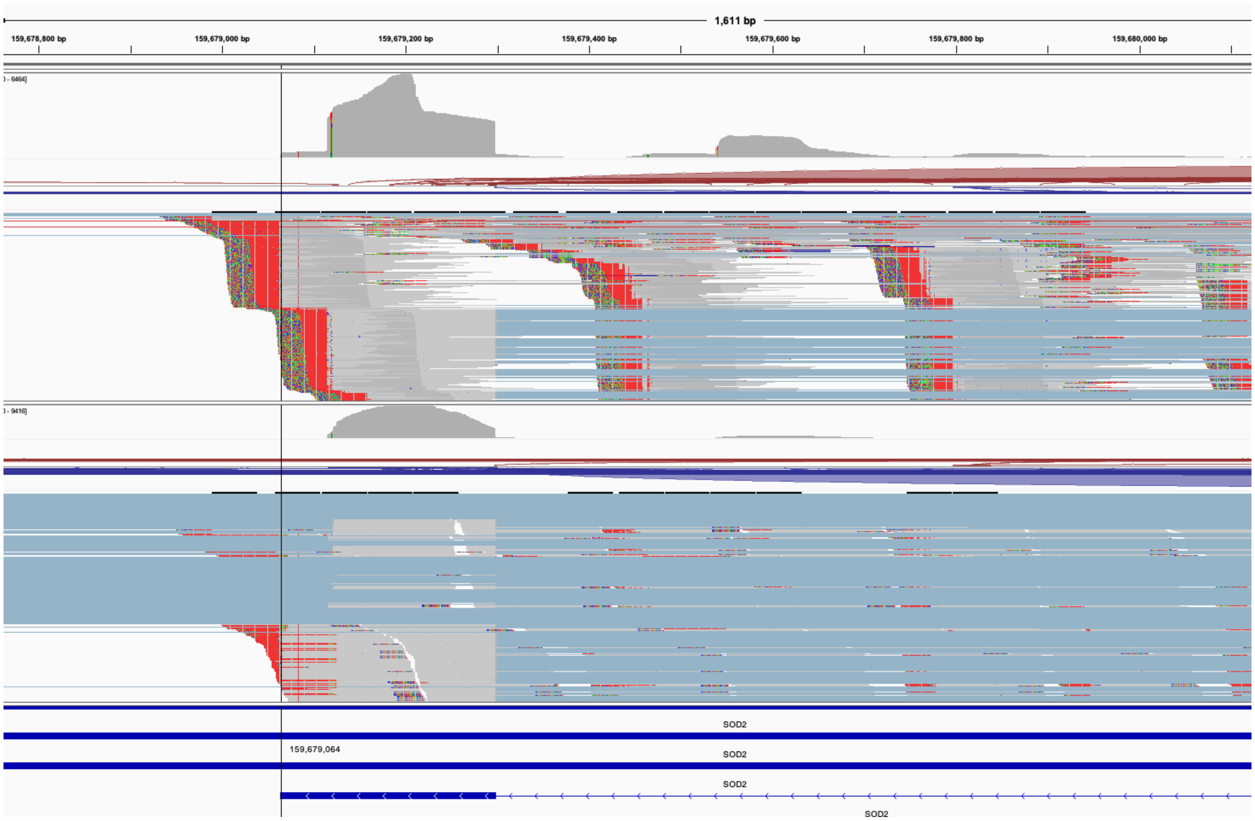


**Supplementary Figure S2**. Paired and single-end read coverage distribution at SOD2. Paired alignments and coverage (top tracks) show evidence of two polyadenylation sites in close proximity, with most reads aligning at the more proximal (right) position. Single-end alignments also show evidence of both positions as soft-clipped tails, but the resulting output groups all alignments into a peak ending at the distal site.

**Supplementary Table 1**

| | | Library S1 | S2 | S3 | S4 | S5 |
|---|---|---|---|---|---|---|
| | SRA accession | SRX7528394 | SRX20356215 | SRX20356216 | SRX20356222 | SRX20356230 |
| | Pearson's R (gene abundance, Element ~ Illumina) | .92 | .99 | .99 | .99 | .99 |
| UMI counts/ cell | Illumina | 14153 | 17169 | 19804 | 13881 | 12939 |
| | Aviti | 13331 | 14833 | 17336 | 7334 | 19640 |
| Reads | Illumina | 346436428 | 510019528 | 485844363 | 426841042 | 333288137 |
| | Aviti | 271981610 | 335283933 | 338136830 | 192278636 | 679945763 |
| Sequencing saturation | Illumina | .75 | .66 | .66 | .20 | .30 |
| | Aviti | .59 | .53 | .57 | .09 | .49 |
| Insert size distr. | Aviti | 266 | 150 | 157 | 160 | 210 |

**Table S1**: Sequencing metrics and QC data. Pearson's R refers to correlation between pseudobulked gene expression estimates between instruments (log of average total counts per gene, for genes with at least 1 count in both samples). Counts were quantified on pre-trimmed R2 alignment data using 'Gene' mode in STARsolo. Sequencing saturation is an estimation of the chance of observing a previously observed UMI with a new read. Median insert size distribution is calculated from paired-end alignments for Aviti data only.

| | **Paired-end Unique \| Multi Aln %** | | **Read 2 Unique \| Multi Aln %** | | **Read 1 Unique \| Multi Aln %** | |
|---|---|---|---|---|---|---|
| Element S1 | 94.1 | 3.2 | 93.2 | 4.8 | 89.5 | 9.3 |
| Element S2 | 88.8 | 6.9 | 87.7 | 7.8 | 83.6 | 12.0 |
| Element S3 | 91.9 | 4.9 | 90.8 | 5.7 | 87.3 | 9.6 |

| | | | | | | |
|---|---|---|---|---|---|---|
| Element S4 | 87.5 | 5.7 | 86.9 | 6.4 | 84.4 | 9.4 |
| Element S5 | 91.3 | 4.8 | 90.7 | 6.0 | 87.1 | 9.6 |
| Illumina S1 | NA | | 91.1 | 4.3 | NA | |
| Illumina S2 | NA | | 81.2 | 8.1 | NA | |
| Illumina S3 | NA | | 86.5 | 5.9 | NA | |
| Illumina S4 | NA | | 81.7 | 6.6 | NA | |
| Illumina S5 | NA | | 85.4 | 6.2 | NA | |

**Table S2**: Unique and multiple read alignment % for paired-end and single-end alignment modes, after implementing pre-trimming step. Pre-trimming step was not applied to Illumina paired and R1 alignment because it does not address the underlying homopolymer error problem, and alignment rates remain very low (<30%).

| **Name** | **Medium** | **Year** | **DOI** |
|---|---|---|---|
| SCAPE-APA | Preprint | 2024 | https://doi.org/10.1101/2024.03.12.584547 |
| Infernape | Published | 2023 | https://doi.org/10.1101/gr.277864.123 |
| SCINPAS | Published | 2023 | https://doi.org/10.1093/nargab/lqad079 |
| scMAPA | Published | 2022 | https://doi.org/10.1093/gigascience/giac033 |
| **scraps** | Preprint | 2022 | https://doi.org/10.1101/2022.08.22.504859 |
| ‘Agarwal et al’ | Published | 2021 | https://doi.org/10.1101/2021.01.21.427498 |
| scDaPars | Published | 2021 | https://doi.org/10.1101/gr.271346.120 |
| scLAPA | Preprint | 2021 | https://doi.org/10.1101/2021.01.04.425335 |
| SCAPTURE | Published | 2021 | https://doi.org/10.1186/s13059-021-02437-5 |
| MAAPER | Published | 2021 | https://doi.org/10.1186/s13059-021-02429-5 |
| scAPAtrap | Published | 2021 | https://doi.org/10.1093/bib/bbaa273 |
| movAPA | Published | 2021 | https://doi.org/10.1093/bioinformatics/btaa997 |

| | | | |
|---|---|---|---|
| APA-seq | Published | 2020 | https://doi.org/10.1093/nar/gkaa359 |
| Sierra | Published | 2020 | https://doi.org/10.1186/s13059-020-02071-7 |
| scDAPA | Published | 2020 | https://doi.org/10.1093/bioinformatics/btz701 |
| scAPA | Published | 2019 | https://doi.org/10.1093/nar/gkz781 |
| **polyApipe** | Poster | 2019 | https://doi.org/10.7490/f1000research.1117076.1 |

**Table S3**: Non-exhaustive summary of existing bioinformatics tools for measuring polyadenylation site usage in 3’ single-cell RNA-seq data. Methods used in this study are in bold.
